# Integrated cancer cell-specific single-cell RNA-seq datasets of immune checkpoint blockade-treated patients

**DOI:** 10.1101/2024.01.17.576110

**Authors:** Mahnoor N. Gondal, Marcin Cieslik, Arul M. Chinnaiyan

**Affiliations:** Department of Computational Medicine & Bioinformatics, University of Michigan, Ann Arbor, MI USA; Michigan Center for Translational Pathology, University of Michigan, Ann Arbor, MI USA; Department of Pathology, University of Michigan, Ann Arbor, MI USA; Department of Urology, University of Michigan, Ann Arbor, MI USA; Howard Hughes Medical Institute, Ann Arbor, MI USA; University of Michigan Rogel Cancer Center, Ann Arbor, MI USA

## Abstract

Immune checkpoint blockade (ICB) therapies have emerged as a promising avenue for the treatment of various cancers. Despite their success, the efficacy of these treatments is variable across patients and cancer types. Numerous single-cell RNA-sequencing (scRNA-seq) studies have been conducted to unravel cell-specific responses to ICB treatment. However, these studies are limited in their sample sizes and require advanced coding skills for exploration. Here, we have compiled eight scRNA-seq datasets from nine cancer types, encompassing 174 patients, and 90,270 cancer cells. This compilation forms a unique resource tailored for investigating how cancer cells respond to ICB treatment across cancer types. We meticulously curated, quality-checked, pre-processed, and analyzed the data, ensuring easy access for researchers. Moreover, we designed a user-friendly interface for seamless exploration. By sharing the code and data for creating these interfaces, we aim to assist fellow researchers. These resources offer valuable support to those interested in leveraging and exploring single-cell datasets across diverse cancer types, facilitating a comprehensive understanding of ICB responses.

## Background & Summary

The cytotoxic activity of T cells can be deactivated through interaction between checkpoint proteins such as PD-1 found on the surface of T cells, and their counterparts, like PD-L1 or PD-L1, present on cancerous cells ^1^. As a result, many cancer cells hijack this mechanism by overexpressing ligands such as PD-L1, disrupting the body’s immune response and impeding the T cells’ ability to eradicate cancer cells ^1^. Research into understanding cancer immune evasion led to the development of immune checkpoint blockade (ICB) therapy that includes monoclonal antibodies that block checkpoint proteins’ functions, allowing activated cytotoxic T cells to eliminate cancer cells ^2^. While such therapies have shown promise in treating different cancer types, their efficacy varies among individual patients and specific cancer types ^3–9^. Numerous studies have attempted to uncover the factors contributing to the success or failure of ICB therapies, predominantly employing RNA expression data ^10–15^. However, due to the multifactorial intrinsic properties of cancer cells, conventional bulk RNA sequencing lacks the resolution needed for an in-depth exploration of cancer cell-specific responses to ICB treatment.

Single-cell RNA-sequencing (scRNA-seq) provides a unique opportunity to dissect the intricate landscape of cancer cell-specific responses to ICB treatment. Numerous scRNA-seq studies have explored patients treated with ICB across various cancer types, including skin cancers such as melanoma ^16–19^, and basal cell carcinoma ^20^. Additionally, investigations have extended to breast cancer subtypes like triple negative ^21^, HER2-positive ^21^, ER-positive ^21^, as well as kidney cancer, specifically clear cell renal carcinoma ^22^, and liver cancers such as hepatocellular carcinoma ^23^, intrahepatic cholangiocarcinoma ^23^. Single-cell studies investigating ICB-treated patients have provided valuable insights into the cellular responses to immune checkpoint blockade therapies, potentially unveiling novel biomarkers. However, these studies often are limited in their sample size and the inherent heterogeneity within tumors and among patients can make it challenging to generalize findings across a broader population. Additionally, single-cell data require sophisticated computational methods due to the large volume of data generated, posing challenges for researchers lacking extensive bioinformatics expertise. Therefore, to overcome limited sample sizes and the complexity of data analysis there is a pressing need to collate, standardize, aggregate, and deposit scRNA-seq datasets with ICB-treated patients to facilitate convenient exploration, ensuring accessibility for both wet and dry lab experts.

Here, we conducted an extensive literature review and meticulously performed data curation, quality control, pre-processing, and analysis. This effort resulted in a robust data resource comprising eight scRNA-seq datasets spanning nine distinct cancer types, involving 174 patients, and 90,270 cancer cells. To enhance accessibility, we have developed user-friendly interfaces housing downsampled single-cell data and pseudobulk data. To create standardized data across various studies, we employed normalization techniques using a gene signature known as stably expressed genes (hSEGs) ^24^. Through our efforts, users can effortlessly explore pseudobulk and standardized datasets via https://scrnaseqicb.shinyapps.io/icbsc_pseudobulk_v12/. This effort provides an unprecedented data resource tailored for investigating ICB responses in cancer cells across a wide range of cancer malignancies. Notably, our resource has been utilized in recent study ^25^ to explore *PIKFYve* gene expressions in responders and non-responders within single-cell melanoma datasets, underscoring the versatility of our data in advancing research on ICB responses in various cancer types. In summary, these resources serve as a support for those keen on leveraging and delving into cancer cell-specific exploration across diverse cancer types, fostering a comprehensive exploration of ICB responses.

## Methods

### Ethical statement

The samples employed in this study were obtained following ethical guidelines and approved by the Institutional Review Board (IRB) in accordance with the protocols established for each respective study. The research strictly adhered to ethical standards and principles governing human subject research.

### Workflow overview

To construct a comprehensive data repository of single-cell RNA sequencing data (scRNA-seq) aimed at investigating cancer cell behavior under ICB treatment, we systematically executed seven key steps, elaborated upon in the following sections (**Fig. 1**).

**Fig.1.**
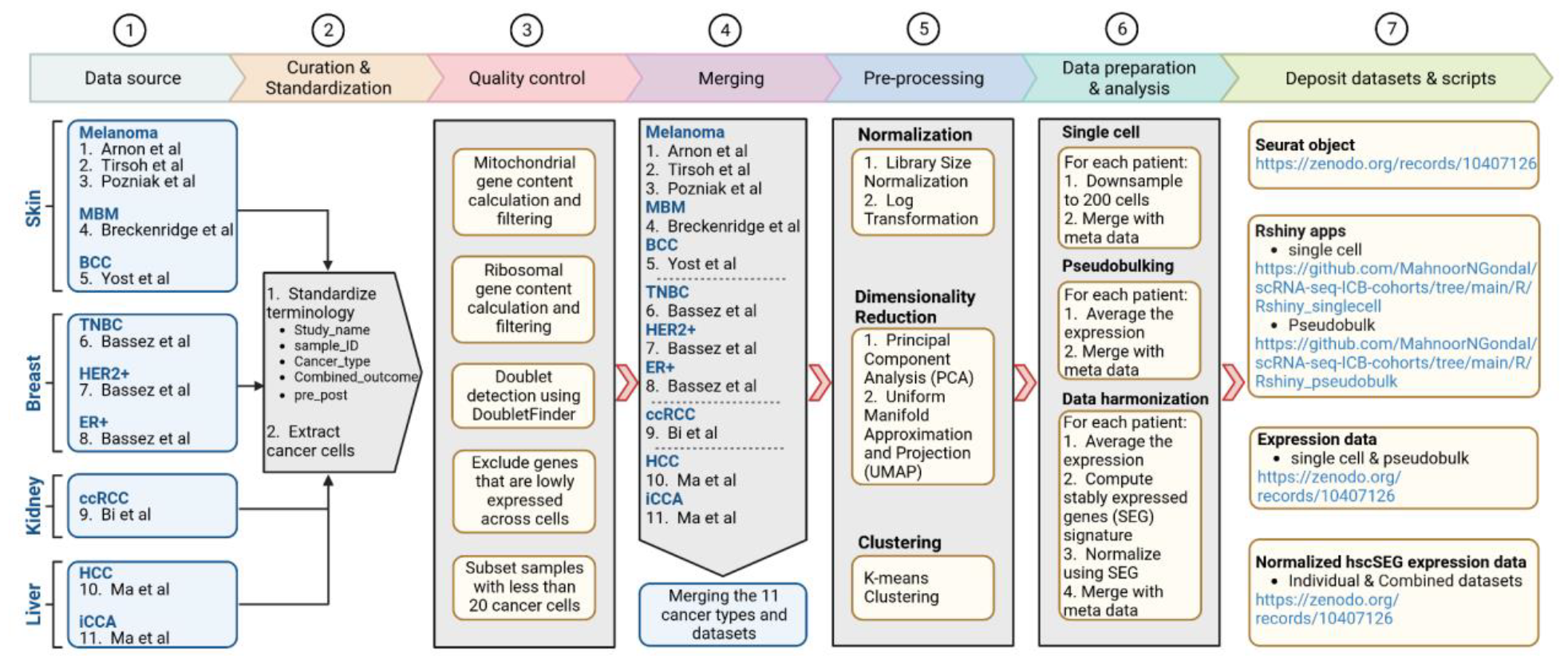
Workflow for developing a dedicated resource centered on cancer cells in ICB studies using single-cell data. The steps include conducting a comprehensive literature review of scRNA-seq studies, standardizing dataset terminologies, extracting cancer cells, implementing rigorous quality control and pre-processing measures, merging datasets for subsequent analysis, and depositing resultant datasets alongside associated R scripts to ensure easy accessibility.

### Data sources

To compile scRNA-seq datasets that detail cancer cell annotations in the context of ICB treatment, an extensive literature survey was undertaken. This survey identified eight key studies, specifically: Jerby-Arnon et al. ^16^, Tirsoh et al. ^17^, Pozniak et al. ^19^, Yost et al. ^20^, Alvarez-Breckenridge et al. ^18^, Bassez et al. ^21^, Bi et al. ^22^, Ma et al. ^23^. These studies encompassed various cancer types, including skin cancers such as melanoma ^16,17,19^, basal cell carcinoma ^20^, melanoma brain metastases ^18^, as well as breast cancer, subtypes triple negative ^21^, HER2-positive ^21^, ER-positive ^21^. Additionally, the studies covered kidney cancer (clear cell renal carcinoma ^22^), and liver cancers like hepatocellular carcinoma ^23^, and intrahepatic cholangiocarcinoma ^23^. The information was extracted from diverse data resources and websites. A comprehensive breakdown of the original data sources and sample specifics is provided in **Table 1**.

**Table 1.** Details of data sources and information. This table presents metadata from eight studies, detailing original publication information, the count of cancer or malignant cells, the number of patients involved, pre-and post-treatment statuses, original data sources, and the locations of processed files.

| # | PMID | Authors | Technology | Cancer type | Cancer subtype | # of malignant cells | # of patients or samples | Pre/UT | Post | Pre/Post (same p) | Primary/ Meta | Location of original data |
| --- | --- | --- | --- | --- | --- | --- | --- | --- | --- | --- | --- | --- |
| 1 | 30388455 | Jerby-Amon et al | SmartSeq2 | Skin | Melanoma | 665 | 8 | 3 (UT) | 5 (Resis) | No | Both | [1-2] |
| 2 | 27124452 | Tirsoh et al | SmartSeq2 |  |  | 602 | 7 | 5 (UT) | 2 (Resis) | No | Met |  |
| 3 | 10.1101/2022.08.11.502598 | Pozniak et al | 10X |  |  | 9201 | 28 | 17 (6 R, 11 NR) | 11 (2 R, 9 NR) | No | Both | [3] |
| 4 | 35706413 | Alvarez-Breckenridge et al | SmartSeq2 |  | MBM | 4990 | 24 | 8 (1 R, 2 NR, 5 UT) | 16 (10 PR, 6 NR) | No | Met | [4] |
| 5 | 31359002 | Yost et al | 10X |  | BCC | 3452 | 8 | 4 [UT] | 4 [2R, 2 NR] | No | Met | [5] |
| 6 | 31588021 | Lichun Ma et al | 10X | Liver | HCC | 743 | 6 | 1 | 5 | No | Both | [6] |
| 7 |  |  |  |  | iCCA | 1025 | 8 | 1 | 7 | No | Both |  |
| 8 | 33958794 | Ayse Bassez et al | 10X | Breast | TNBC | 32501 | 36 | 18 [8 E, 9 NE, 1 n/a] | 18 [8 E, 9 NE, 1 n/a] | Yes | Pri | [7] |
| 9 |  |  |  |  | HER2 | 3108 | 8 | 4 [1 E, 2 NE, 1 n/a] | 4 [1 E, 2 NE, 1 n/a] | Yes | Pri |  |
| 10 |  |  |  |  | ER+ | 27927 | 35 | 18 [3 E, 15 NE] | 17 [2 E, 15 NE] | Yes | Pi |  |
| 11 | 33711272 | Bi et al | 10X | Kidney | ccRCC | 6056 | 6 | 3 [UT] | 3 [2 PR, 1 SD] | No | Met | [8] |

[1-2] - Jerby-Amon + Tirsoh - https://www.ncbi.nlm.nih.gov/geo/query/acc.cgi?acc=GSE115978
[3] - Pozniak - https://rdrr.kuleuven.be/dataset.xhtml?persistentId=doi:10.48804/GSAXBN
[4] - Alvarez-Breckenridge - https://singlecell.broadinstitute.org/single_cell/study/SCP1493/microenvironmental-correlates-of-immune-checkpoint-inhibitor-response-in-human-melanoma-brain-metastases-revealed-by-t-cell-receptor-and-single-cell-ma-sequencing#study-download
[5] - Yost - https://www.ncbi.nlm.nih.gov/geo/query/acc.cgi?acc=GSE123813
[6] - Ma - https://www.ncbi.nlm.nih.gov/geo/query/acc.cgi?acc=GSE125449
[7] - Bassez - https://lambrechtslab.sites.vib.be/en/single-cell
[8] - Bi - https://singlecell.broadinstitute.org/single_cell/study/SCP1288/tumor-and-immune-reprogramming-during-immunotherapy-in-advanced-renal-cell-carcinoma#study-download

### Data curation and standardization

We performed meticulous data extraction, curation, and standardization for each individual single-cell study, drawing from diverse resources and databases. Acknowledging potential discrepancies arising from varying naming conventions and cell annotations among different authors, we anticipated discrepancies. To maintain uniformity and coherence within our resources, we took steps to standardize terminologies across all studies before proceeding with the extraction of cancer cells (**Fig. 2**). Additional specifics for each study are provided below for clarity.

**Fig.2.**
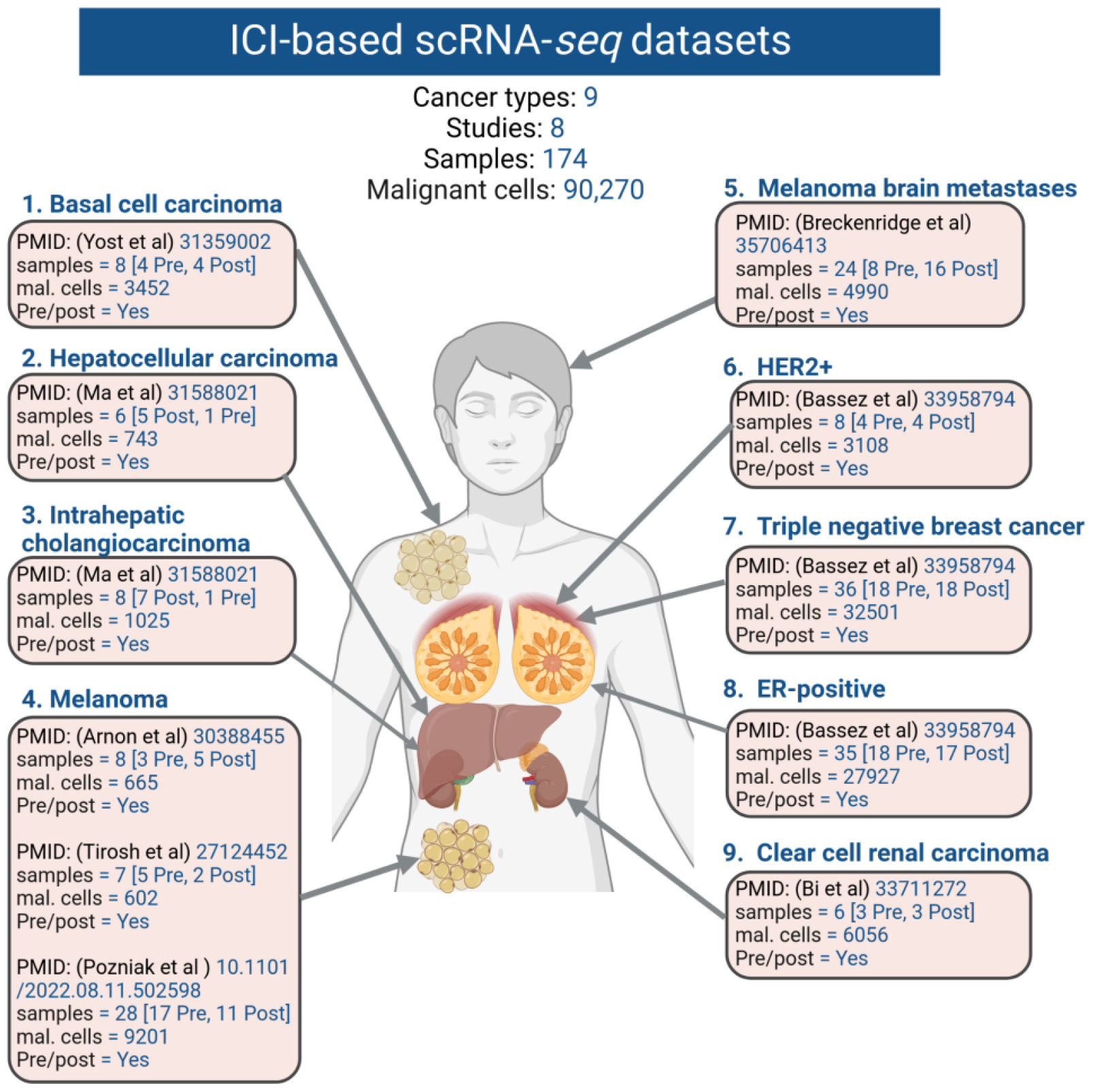
Overview of single-cell studies investigated in this resource. The basal cell carcinoma dataset comprised 3452 cancer cells, while the hepatocellular carcinoma dataset contained 743 cancer cells, and the intrahepatic cholangiocarcinoma dataset included 1025 cells. The combined three melanoma datasets yielded a total of 10,468 malignant cells, with the melanoma brain metastasis dataset contributing 4990 cells. Within the breast cancer datasets (HER2+, TNBC, ER+), there were 3108, 32,501, and 27,927 cancer cells, respectively. Additionally, the clear cell carcinoma dataset comprised 6056 cells. In total, the resulting dataset amalgamates findings from eight studies encompassing nine distinct cancer types, encapsulating a sum of 90,270 malignant cells.

### Jerby-Arnon et al. and Tirsoh et al. (Melanoma)

Jerby-Arnon et al. ^16^ unveiled a cancer cell mechanism fostering T cell exclusion, contributing to resistance against checkpoint blockade therapies. This program elucidated the mechanisms behind the evasion of immune surveillance in tumors, impacting the effectiveness of immunotherapy. In contrast, Tirosh et al. ^17^ leveraged scRNA-seq to unravel the complex multicellular landscape of metastatic melanoma. It comprehensively profiled individual cells within the tumor microenvironment, offering insights into cellular interactions, heterogeneity, and potential therapeutic targets for this aggressive cancer.

The Jerby-Arnon datasets for melanoma samples were extracted from GSE115978. The study builds upon the previously published melanoma study by Tirsoh et al. and as a result, already contained the previous samples. Therefore we standardized both Jerby-Arnon et al. and Tirosh et al.’s melanoma studies together. The metadata was also supplied from the Tumor Immune Single Cell Hub (TISCH) ^26^. The data was first converted to a Seurat object using raw counts data, and cell annotation files. The samples were renamed to “sample_id”. As mentioned in the original publication, all treated patients were classified as resistant to ICB treatment and categorized as “Unfavourable” in the “Combined_outcome” criteria. InferCNV was employed to identify malignant cells in both cohorts. Access to the processed Seurat object in the RDS file and the associated Rmd file for this dataset has been provided for accessibility (**Table 2**).

**Table 2.**
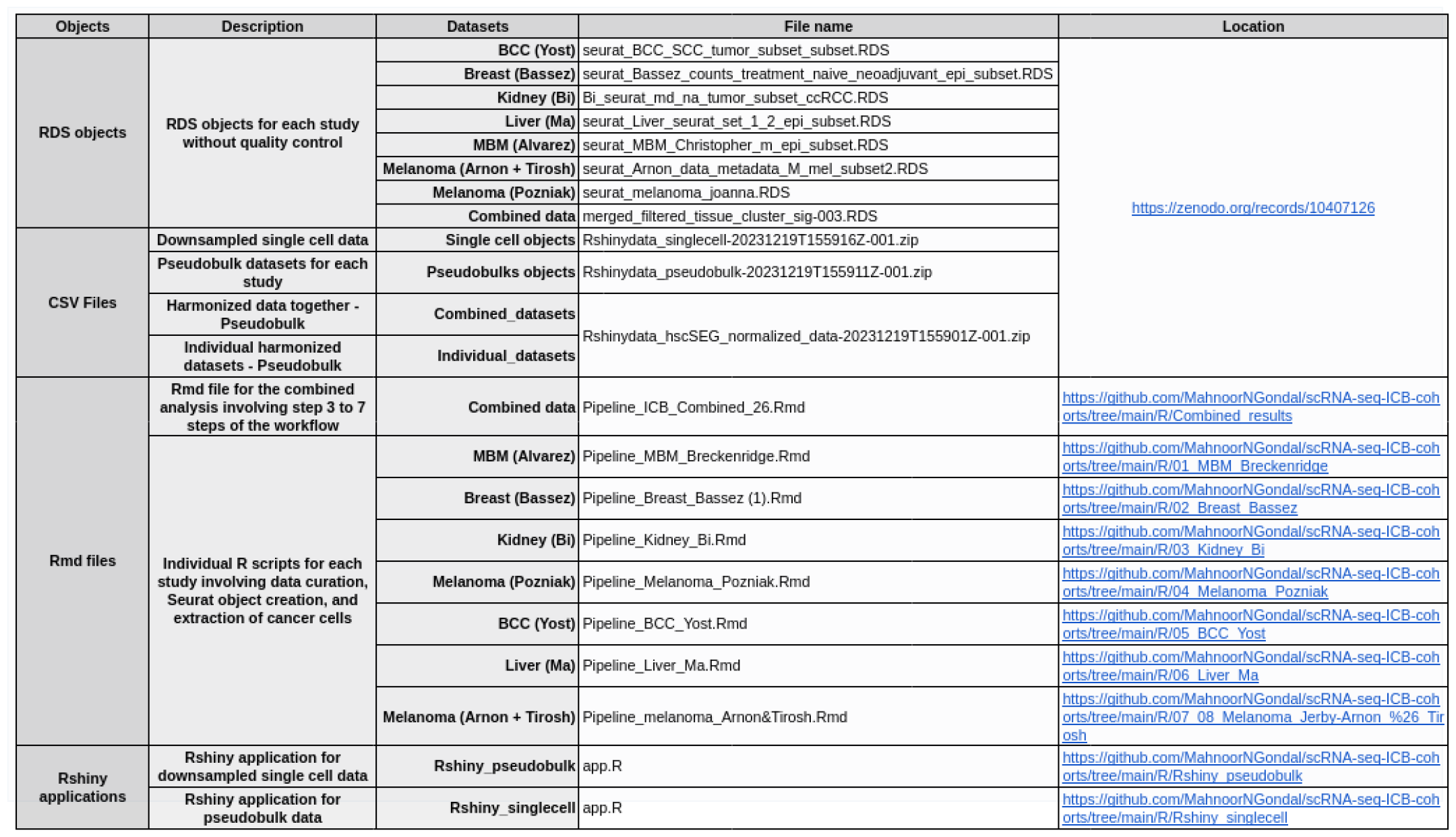
Summary of code and data-related Items. This table offers a comprehensive overview, encompassing file types, detailed descriptions, names, and locations. Its purpose is to streamline accessibility and serve as a convenient point of reference for users seeking access to code and data resources.

### Pozniak et al. (Melanoma)

A recently published melanoma scRNA-seq study by Pozniak et al. ^19^ was utilized in our resource. This study highlighted the significant impact of intrinsic resistance on ICB effectiveness in melanoma. By analyzing scRNA-seq data from patients undergoing ICB therapy, the study identified TCF4 as a key regulator of the Mesenchymal-like (MES) program, suggesting a potential therapeutic avenue to enhance immunogenicity in ICB-resistant melanomas and improve sensitivity to targeted therapy.

The dataset from this study was accessible through KU Leuven Research Data Repository (RDR) in the form of RDS files with metadata included. This was already a Seurat object with raw count data included as well as the clinical information for both responders and non-responders. Responders were categorized as “Favourable” and non-responders as “Unfavourable” in the “Combined_outcome” criteria. Inferred CNV profiles based on scRNA-seq data were employed to identify malignant cells using the HoneyBADGER method ^27^. The processed Seurat object in the RDS file and the associated Rmd file for this dataset are made accessible (**Table 2**).

### Yost et al. (Basal Cell Carcinoma)

Yost et al. ^20^ study, revealed the phenomenon of clonal replacement where new tumor-specific T cell clones emerged after PD-1 blockade treatment in Basal Cell Carcinoma (BCC). This replacement suggested a dynamic shift in the T cell population, potentially impacting the efficacy of PD-1 blockade therapy in the past.

scRNA-seq data for this study was available on GSE123814, alongside metadata information. Raw counts and metadata were employed to create a Seurat object. The dataset also contained squamous cell carcinoma (SCC) samples, however, since SCC lacked malignant cells we removed it from the final dataset. Single-cell CNVs were detected to label malignant cells using HoneyBADGER ^27^. Those with outcomes “Yes” were labeled as “Favourable” and those with “No” were categorized as “Unfavorable” in the “Combined_outcome” criteria. The processed Seurat object in the RDS file, along with the associated Rmd file for this dataset, is now accessible (**Table 2**).

### Alvarez-Breckenridge et al. (Melanoma Brain Metastasis)

Melanoma brain metastases (MBM) refer to the spread of malignant melanoma cells from their primary site to the brain, forming secondary tumors. These metastases present significant challenges due to their aggressive nature, often leading to severe neurological complications and impacting treatment options. Alvarez-Brechenridge et al.^18^ explored the microenvironmental changes in human melanoma brain metastases triggered by immune checkpoint inhibition. The study provided insights into the alterations within the brain metastatic sites in response to immune checkpoint therapy, shedding light on the intricate interplay between the tumor microenvironment and treatment efficacy.

This dataset was accessible using Broad Institute’s single-cell portal. We downloaded the raw count data from the portal alongside the barcode and features matrix to create the Seurat object. The metadata was later added using the AddMetaData function in the Seurat package. Malignant cells were labeled using inferred copy-number profiles within cells, from a single patient, using all FACS gating categories (CD45−, CD45+, CD3+). The resultant copy number profiles were then projected onto their first principal component towards generating a “malignancy score” which was later utilized for malignant cell detection. Responders and partial-responders were categorized as “Favourable” and non-responders as “Unfavourable” in the “Combined_outcome” criteria. The processed Seurat object stored in the RDS file, along with the associated Rmd file for this dataset, is available for access (**Table 2**).

### Bassez et al. (HER2+, ER+, Triple-negative Breast Cancer)

The Bassez et al.^21^ study created a single-cell map, detailing intratumoral alterations occurring during anti-PD1 treatment in breast cancer patients (TNBC, HER2+, ER+). It provided insights into the dynamic changes within the tumor microenvironment in response to anti-PD1 therapy, aiding in understanding treatment effects and potential therapeutic avenues.

This dataset encompassed two cohorts; Cohort 1 contained treatment-naive samples while Cohort 2 had neo-adjuvant therapy samples. Datasets including the two cohorts’ counts and metadata are easily accessible through the VIB-KU Leuven Center for Cancer Biology website at https://lambrechtslab.sites.vib.be/en/single-cell. Both cohorts were first converted to Seurat objects and then combined for downstream analysis. InferCNV was employed to identify copy number alterations in the cells to label malignant cells in both cohorts. The clinical outcomes in this study were divided between “expanded” and “non-expanded”. As such, expanded samples were categorized as “Favourable” and non-expanded as “Unfavourable” in the “Combined_outcome” criteria. The Seurat object, processed and stored in the RDS file, as well as the corresponding Rmd file for this dataset, are both accessible (**Table 2**).

### Bi et al. (Clear Cell Renal Carcinoma)

Bi et al. ^22^ study documented tumor and immune system reprogramming occurring during immunotherapy for renal cell carcinoma patients. It offered insights into the dynamic changes within both the tumor and immune cells, shedding light on the mechanisms involved in immunotherapy response and resistance.

The dataset was downloaded from Curated Cancer Cell Atlas (3CA; https://weizmann.ac.il/sites/3CA) ^28^ database including the raw counts, features, and barcodes alongside patients’ clinical information. These were employed to create a Seurat object. Inferred Copy Number Aberration (infercna) (using the package available at https://github.com/jlaffy/infercna) was utilized to identify malignant cells. Partial-responders were categorized as “Favourable” and stable disease as “Unfavourable” in the “Combined_outcome” criteria. The processed Seurat object, encapsulated within the RDS file, and its related Rmd file for this dataset have been made readily available for access and utilization (**Table 2**).

### Ma et al. (Hepatocellular Carcinoma, Intrahepatic Cholangiocarcinoma)

Ma et al.’s ^23^ research illustrated that the variability within liver cancer cells played a significant role in reshaping the tumor microenvironment. It emphasized how the diversity among cancer cells impacted the changes occurring in the surrounding tumor environment, offering a critical understanding of the intricate relationship between cancer cells and their immediate surroundings.

Both Hepatocellular carcinoma (HCC) and intrahepatic cholangiocarcinoma (iCCA) datasets were downloaded from GSE125449 including raw counts, barcodes, and features to create two Seurat objects. The two Seurat objects were then merged. InferCNV was employed to identify malignant cells in both cohorts. The processed Seurat object stored within the RDS file and the associated Rmd file for this dataset are now accessible for utilization and reference purposes (**Table 2**).

### Quality control

The quality control steps are described in detail under technical validation.

### Merging

For subsequent analysis, we used Seurat’s merge function to combine the eight datasets. The resulting merged data comprises 174 patients and 90,270 cancer cells, spanning nine cancer types.

### Pre-processing

Single-cell analysis was performed using the Seurat package (v4.1.1). To pre-process the datasets we employed Seurat’s standardized pipeline. Data normalization was performed using log1p normalization from Seurat’s NormalizeData function. Dimensionality reduction and clustering were undertaken using an unsupervised graph-based clustering approach using 20 dimensions. Uniform Manifold Approximation and Projection (UMAP) was employed to visualize the data. The UMAP was visualized using individual cancer types across pre and post-samples (**Fig. 3A**). We also observed UMAP in light of ‘Favourable,’ ‘Unfavourable,’ ‘Untreated’ (UT), and ‘n/a,’ combined outcomes across both study and cancer types (**Fig. 3B**). Detailed sample and study information is available in **Table 1**. Furthermore, a comprehensive overview of the datasets, including the number of samples per study, sample sizes in pre and post-datasets, and cancer cells within ‘Combined_outcomes,’ is presented in **Fig. 3C-E**.

**Fig.3.**
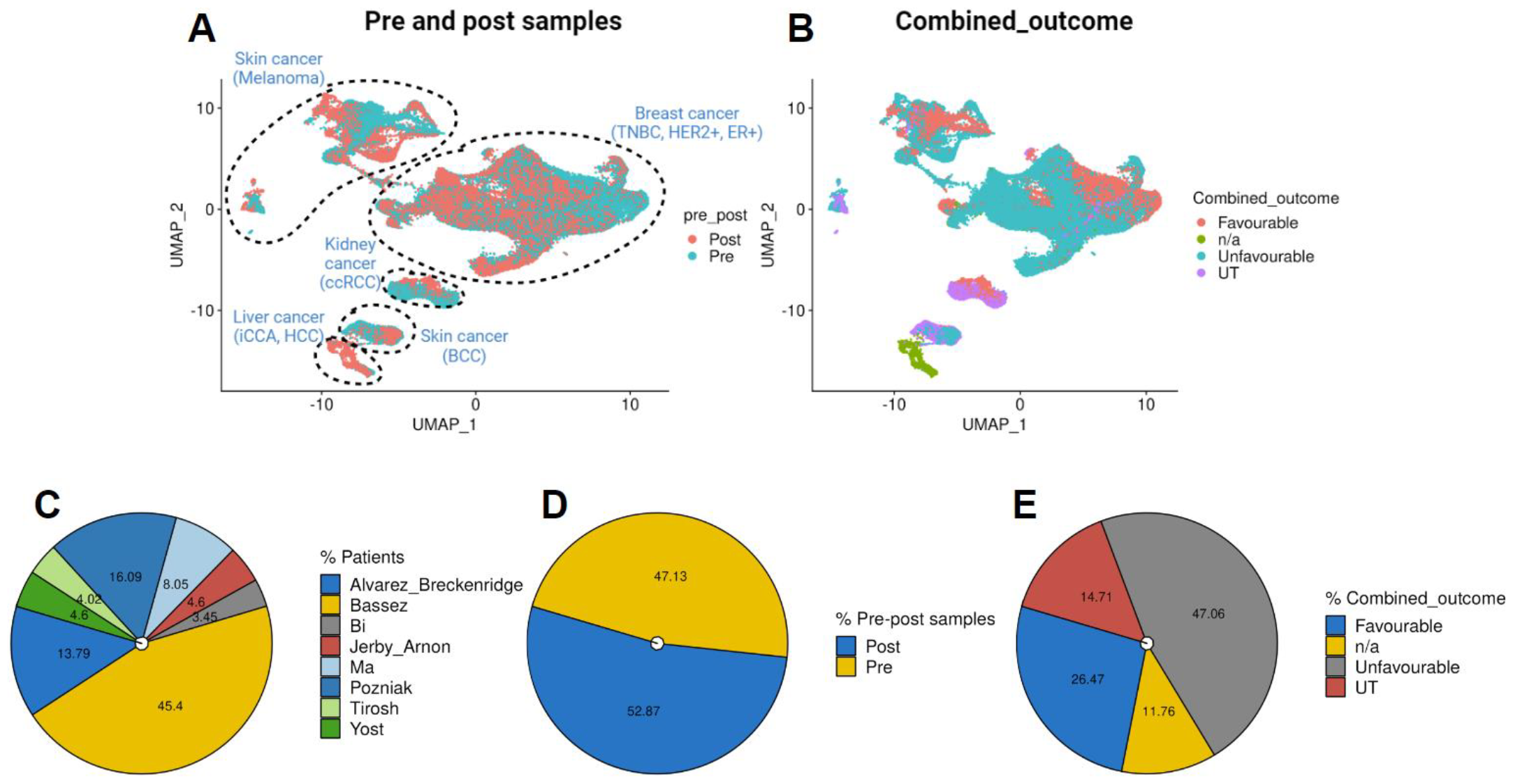
Visualization of data distribution across studies. **(A)** UMAP visualizations depicting pre-and post-treatment status, and cancer type information. **(B)** UMAP for combined outcome distributions within different datasets. **(C-E)** Pie charts illustrating patient distribution across studies, total pre- and post-treatment samples, and ‘Favourable,’ ‘Unfavourable,’ ‘Untreated’ (UT), and ‘n/a,’ combined outcomes across studies in the merged dataset.

### Data preparation and analysis

We have made the Seurat objects available for use in the form of RDS files the details of which are provided in **Table 2**. Analyzing single-cell data demands advanced computational methods owing to the vast amount of generated data, presenting challenges for researchers lacking deep bioinformatics knowledge. To facilitate accessibility for non-bioinformaticians, we deposited CSV files necessary for data exploration as well as developed Rshiny applications. The Rshiny applications are a user-friendly tool that provides access to three distinct data types within the application interface. Further specifics on the data generation process are outlined below.

### Single-cell data

To analyze the single-cell RNAseq datasets within the Rshiny application, we initiated by downsampling 200 cancer cells from each sample in both pre and post-treatment contexts. Following this, we segregated each study and integrated it with its corresponding metadata. These generated files were downloaded as CSV formats and archived on Zenodo and Google Drive. For real-time importation during the Rshiny application runtime, the files were accessed using the googledrive package.

### Pseudobulk data

To manage extensive datasets and circumvent zero values in the raw count for single-cell data, we generated pseudobulks for individual samples in both pre and post-treatment settings. This involved subsetting to each study and employing Seurat’s AverageExpression function to extract the pre and post-cluster expression of all genes. Subsequently, this combined data, integrated with study metadata, was downloaded in CSV format and archived on Zenodo and Google Drive. To enable real-time data importation within the Rshiny application, we utilized the googledrive package to access the files. The data size was modest enough to be effectively utilized within the rsconnect application and is accessible here https://scrnaseqicb.shinyapps.io/icbsc_pseudobulk_v12/.

### Harmonized data

Single-cell cohorts often face limitations due to sample size. To collectively analyze these cohorts, we normalized the datasets by initially segmenting them into each study. Using the AverageExpression function, we created pseudobulks for pre and post-samples. Next, we employed stably expressed genes (hSEGs) gene signature ^24^ to normalize within each cohort, effectively correcting for batch effects. Each dataset was then combined into one and archived on Zenodo and Google Drive in a combined and individual structure. This data can also be explored using the rsconnect application here https://scrnaseqicb.shinyapps.io/icbsc_pseudobulk_v12/.

### Deposit datasets and scripts

Finally, we have prioritized accessibility by ensuring the R script’s availability to the users. We achieved this by archiving the scripts on GitHub, to encourage collaboration, version control, and users to engage with and contribute to the codebase. Additionally, we developed user-friendly Rshiny applications, providing an intuitive interface for efficient data interaction. Furthermore, to enable users to easily access and reference the data, we shared the data in the form of CSV and RDS files on Zenodo, fostering an open and collaborative environment, details provided in **Table 2**.

## Data Records

The data files can be downloaded from the Zenodo data platform in CSV and RDS formats https://zenodo.org/records/10407126. The Rmd R scripts and Rshiny applications are available on GitHub https://github.com/MahnoorNGondal/scRNA-seq-ICB-cohorts. The pseudobulk application is available through rsconnect: https://scrnaseqicb.shinyapps.io/icbsc_pseudobulk_v12/.

The individual files are described in **Table 2**.

## Technical Validation

For each study, a comprehensive series of quality control measures were systematically implemented to ensure the robustness of the single-cell data. The original dataset encompassed a total of 100,025 cancer cells. To safeguard data quality, the first step involved the identification and removal of genes exhibiting high mitochondrial content, which is indicative of potentially compromised or dead cells ^29^. The percentage of mitochondrial gene content was calculated for each cell using the PercentageFeatureSet function. Cells surpassing a 20% threshold in mitochondrial gene content were excluded, resulting in the removal of 4.48% of cells (**Fig. 4A**).

**Fig.4.**
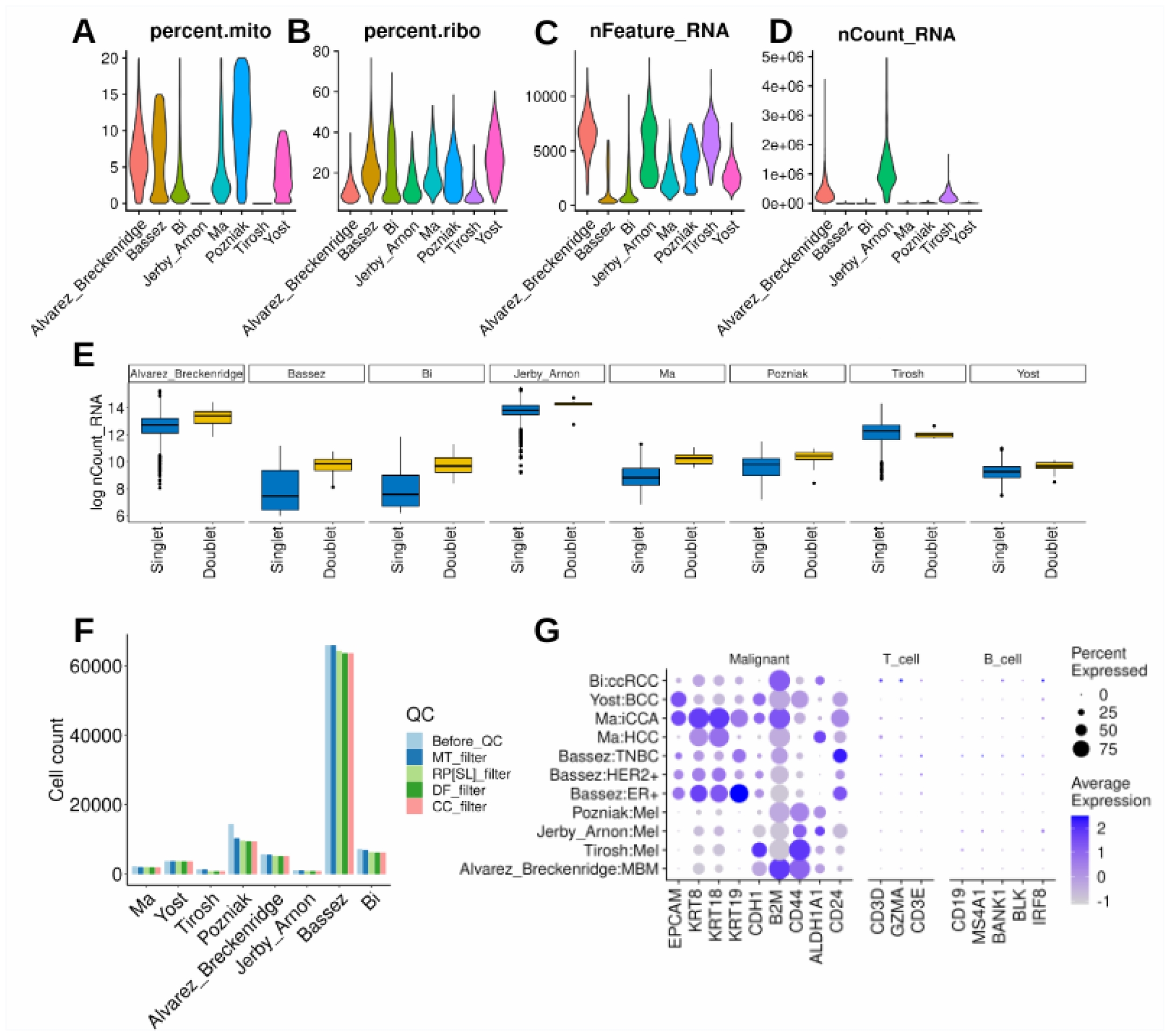
Quality control and technical validation across studies. **(A-D)** Outline the data subsetting process, retaining cells with less than 20% mitochondrial gene content, ensuring genes are expressed in at least 3 cells, and retaining cells with a ribosomal gene content of at least 5%. **(E)** Demonstrates the execution of DoubletFinder, filtering the dataset to retain only ‘Singlet’ cells. **(F)** visually represents the number of cancer cells retained at each filtering step (Before_QC: before quality control, MT_filter: mitochondrial gene content filter, RP[SL]_filter: ribosomal gene content filter, DF_filter: DoubletFinder filter, CC_filter: cancer cells per patient filter). **(G)** showcases highlighted markers specific to epithelial (malignant) cells ensuring data quality.

Subsequently, to enhance the quality of the dataset, an additional criterion was applied, focusing on ribosomal gene content. Cells with less than 5% ribosomal RNA expression were excluded using the same function, leading to the retention of 91,320 cancer cells (**Fig. 4B**). Further refinement involved filtering the dataset to retain only genes expressed in a minimum of 4 cells (**Fig. 4C-D**).

To address potential technical issues arising from doublets ^29^ (instances of more than one cell per bead), the DoubletFinder ^30^ algorithm was employed with default parameters for each study. This process identified and eliminated ‘Doublets,’ or low-quality cells leaving only ‘Singlet’ cells and resulting in 90,407 cancer cells (**Fig. 4E**).

Additional refinement steps included ensuring that each patient’s data contained a minimum of 20 malignant cells. **Fig. 4F** breaks down the number of cells at each step of the quality control process. The final datasets, after these quality control steps, comprised 90,270 cancer cells, reflecting the removal of 9.75% of the initial cells.

Validation of the data was conducted using established markers for malignant (epithelial), T cells, and B cells. Remarkably, the validation process underscored markers exclusively associated with malignant cells (**Fig. 4G**), affirming the quality and specificity of the refined datasets.

## Code Availability

The code can be accessed on GitHub, https://github.com/MahnoorNGondal/scRNA-seq-ICB-cohorts. All packages and their versions are mentioned on GitHub. The datasets can be assessed freely from zenodo, https://zenodo.org/records/10407126.

### Acknowledgments

The study was supported by the National Cancer Institute (NCI) Outstanding Investigator Award R35CA231996 (A.M.C.), NCI Prostate SPORE grant P50CA186786 (A.M.C.), and NCI Michigan-VUMC Biomarker Characterization Center grant U2CCA271854 (A.M.C.). A.M.C. is also a Howard Hughes Medical Institute Investigator, A. Alfred Taubman Scholar, and American Cancer Society Professor. This manuscript was also supported in part by funding from the Innovation in Cancer Informatics (ICI398672) and the V Foundation for Cancer Research (T2019-006) to M.C.

## Author contributions

MC, AMC, and MNG carried out the study design and drafted the manuscript. All authors contributed to the article and approved the submitted version.

## Competing interests

A.M.C. is a co-founder of and serves as a Scientific Advisory Board member for LynxDx, Esanik Therapeutics, Medsyn, and Flamingo Therapeutics. A.M.C. is a scientific advisor or consultant for EdenRoc, Aurigene Oncology, Ascentage Pharma, Proteovant, Belharra, Rappta Therapeutics, and Tempus.

## Notes

https://zenodo.org/records/10407126

https://github.com/MahnoorNGondal/scRNA-seq-ICB-cohorts

### Summary of Updates

We have revised this version to update the "Acknowledgments" section and rearranged the figures/tables only.

